# A simplified electronic circuit for combined single-cell stimulation and recording using loose cell-attached electrodes

**DOI:** 10.1101/191718

**Authors:** Ben W. Strowbridge, R. Todd Pressler

## Abstract

While tight-seal patch clamp recordings have found wide use in neuroscience and in other fields, the requirement to replace the glass pipette after every attempted recording represents an impediment to high throughput studies such as searching for monosynaptically connected pairs of neurons. Loose cell-attached recording was introduced in 2000 to circumvent this problem since it enabled combined recording and stimulation of visually-identified neurons without necessitating a tight (gigaohm) seal. Since the stimulus voltages required to evoke action potentials through low resistance seals are beyond the capacity of most commercial amplifiers, Barbour and Isope introduced a variation of classic patch clamp amplifier circuit that is able to deliver stimulus voltages that are effective in triggering action potentials under the loose cell-attached patch clamp configuration. The present report presents the design and operation of a simpler amplifier that contains only two integrated circuits and is able to effectively stimulate and record action potentials in mitral cells in rodent olfactory bulb slices. The addition of an accessory analog gating circuit enables manual control of the stimulus voltage with pulse timing controlled by a digital output from a computer. This system may be useful in studies that require surveying many potential pairs of neurons for synaptic connections and for sampling and manipulating single-cell activity in in vivo electrophysiology experiments.

## Introduction

Patch clamp methods have had a major impact in cellular neuroscience and have facilitated recordings of activity at both single channel (**10**) and whole cell (**7**) recording modes.Because of its high sensitivity and low noise, whole-cell recording has proved especially useful in probing synaptic circuits in acute and cultured brain slices. However there are two primary disadvantages associated with intracellular recordings using the whole-cell recording configuration: dialysis of the intracellular contents of the recorded neuron and the requirement to replace the recording pipette after every recording attempt. In 2000, Barbour and Isope (**4**) introduced a novel recording configuration for brain slice recording, loose cell-attached recording (LCA), that circumvents both problems and a provides a convenient means for both triggering and recording spiking in individual neurons. This method is a variation of traditional cell-attached recording (**8**, **9**, **15**, **19**) that had been used many groups to monitor neural spiking (e.g., **5**, **6**, **12**). Loose cell-attached recording was devel-oped originally for recording from muscle cells (**1**, **22**). With LCA, the same pipette can be used to record from multiple neurons since the seal resistance required to record spiking is less than needed to detect channel activity. Since there is no whole-cell access with LCA,there is also no intracellular dialysis. The ability to patch multiple neurons using the same pipette is especially useful for locating synaptically-coupled pairs of neurons (**5**, **16**).

One reason why LCA was not commonly used is that the voltages required to trigger APs (typically 100s of mV) are beyond the capacity of most commonly used patch clamp amplifiers. Blindly stimulating at these voltages without recording the neuronal response is not generally a useful strategy since the effective stimulus intensities can damage the neuron. To be useful, it is necessary to know whether the stimulus applied was effective in triggering an action potential (AP) and then employ the smallest stimulus intensities possible. The primary innovation in the Barbour and Isope (**4**) study was the introduction of a specialized patch clamp circuit that allowed relatively high voltages to be applied to the membrane patch in the LCA configuration while retaining the ability to detect spiking. With this type of patch clamp amplifier it was often possible to resolve an evoked AP beyond the large stimulus artifact in the recording channel. Even if APs could not be discerned in the on-line recording, evoked spikes could be easily resolved by subtracting the stimulus artifact associated with a failure trial or a just-subthreshold response. The ability to both stimulate and detect APs through LCA recording represented an advance over blindly stimulating through a LCA recording without verification that the stimulus had triggered an AP.

While the Barbour and Isope circuit was successful at performing LCA recording, it was moderately complex, containing 8 operational amplifiers (op amps) in separate headstage and remote electronic circuits. The design also included additional features, such as bath ground isolation, that add to circuit complexity. In this report we describe a simpler design that contains only a headstage circuit and requires two op amps. We show this simpler circuit is effective in both recording and stimulating APs from mitral cells in acute olfactory bulb slices. The printed circuit board design for the headstage amplifier is freely available through the GitHub repository (*https://github.com/StrowbridgeLab/LPA*). We also describe an analog stimulation control device that provides an alternative to specifying stimulus intensities using software interface controls.

## Methods

### Slice Preparation

Horizontal olfactory bulb slices 300 *μ*m thick were made from ketamine-anesthetized P14-25 Sprague-Dawley rats of both sexes as previously described (**2**, **3**, **11**). Slices were incubated for 30 min at 30°C and then at room temperature until use. All experiments were carried out in accordance with the guidelines approved by the Case Western Reserve University Animal Care and Use Committee. Slices were placed in a recording chamber and superfused with oxygenated artificial cerebrospinal fluid (ACSF) at a rate of 1.5 ml/min. Recordings were made between 29-32°C. ACSF consisted of (in mM): 124 NaCl, 3 KCl, 1.23 NaH_2_PO_4_, 1.2 MgSO_4_, 26 NaHCO_3_, 10 dextrose, 2.5 CaCl_2_, equilibrated with 95% O_2_/ 5% CO_2_. All drugs were added to the submerged recording chamber by changing the external solution source.

### Whole-Cell Recordings

All whole-cell patch-clamp recordings were made with Axopatch 1C or 1D amplifiers (Axon Instruments) using borosilicate glass pipettes (TW150F-4, WPI) of impedances ranging from 2-5 mOhmx pulled on a P-97 pipette puller (Sutter Instruments). Under whole-cell (WC) current-clamp conditions, recording electrodes contained (in mM): 140 K-methylsulfate (MP Biochemicals), 4 NaCl, 10 HEPES, 0.2 EGTA, 4 MgATP, 0.3 Na_3_GTP, 10 phosphocreatine(pH 7.3 and ∼290 mOsm). Whole-cell recordings were low-pass filtered at 5 kHz (FLA-01,Cygus Technology) and digitized at 10-40 kHz using an ITC-18 simultaneously-sampling data acquisition interface using custom software written in Visual Basic (Microsoft). While the most rapid acquisition rates were not required to sample the low-pass filtered intracellular AP response, faster acquisition speeds were useful for generating the rapid TTL output pulses (25 - 50 *μ*s duration) for controlling the analog switch in the LCA stimulation circuit(see Results below for details).

### DIC Imaging

Slices were imaged using infrared differential interference contrast (IR-DIC) optics on Zeiss Axioskop FS1 or Olympus BX51WI upright microscopes. Transmitted light was restricted to 710-790 nm using a band-pass interference filter placed above the microscope field stop. DIC images were captured using a frame-transfer CCD camera (Cohu) and displayed on a high-resolution monochrome analog monitor (Sony). Individual neurons were visualized using IR-DIC video microscopy before attempting either WC or LCA recording. Neuronal cell type was determined based on IR-DIC morphology and soma laminar location.

### Suppliers

Printed circuit board: Bay Area Circuits, Fremont CA (https://bayareacircuits.com/). Printed circuit design software: DipTrace (http://www.diptrace.com/). Electronic parts and Pomana cases: Digikey, Thief River Falls, MN (https://www.digikey.com/). Electrode holder: A-M Systems, Carlsborg, WA (https://www.a-msystems.com/). Electrode glass: World Precision Instruments, Sarasota, FL (https://www.wpiinc.com/). Electronic filter: Cygnus Technology, Delaware Water Gap, PA (http://www.cygnustech.com/).

## Results

### Headstage amplifier circuit

The core circuit we employed is shown in Fig. 1A and is designed to record APs through the loose patch clamp recording configuration. The headstage circuit (Fig. 1A) was adapted from reference **4**. Both the first-stage FET input op amp (U1, OPA604, Burr Brown/TI) and the differential output amplifier (U2, INA106, Burr Brown/TI) were mounted on a custom printed circuit board manufactured by Bay Area Circuits and enclosed within a small aluminum box connected to the circuit ground. The circuit ground was connected to the same silver chloride-coated silver bath electrode as the whole cell amplifier. A 20 MΩ feedback resistor (Rf; Slim-Mox; Ohmite) set the transimpedance gain. While many high gain tran-simpedance amplifiers include a small capacitor across the feedback resistor to minimize gain peaking, we found that eliminating this extra 1-2 pF capacitance improved our ability to recognize evoked APs even without artifact subtraction. (Holes for a small capacitor across Rf are included in the printed circuit board layout but were not populated.) The± 15 V power to each op amp was bypassed to ground using 0.1 *μ*F ceramic capacitors (not included in the schematic diagram in Fig. 1 but included in the printed circuit board). Through the combination of the input stage (OPA604 with Rf) and the 10x differential amplifier (INA106), the headstage output voltage reflects a scaling of 5 nA/V.

**Figure 1:**
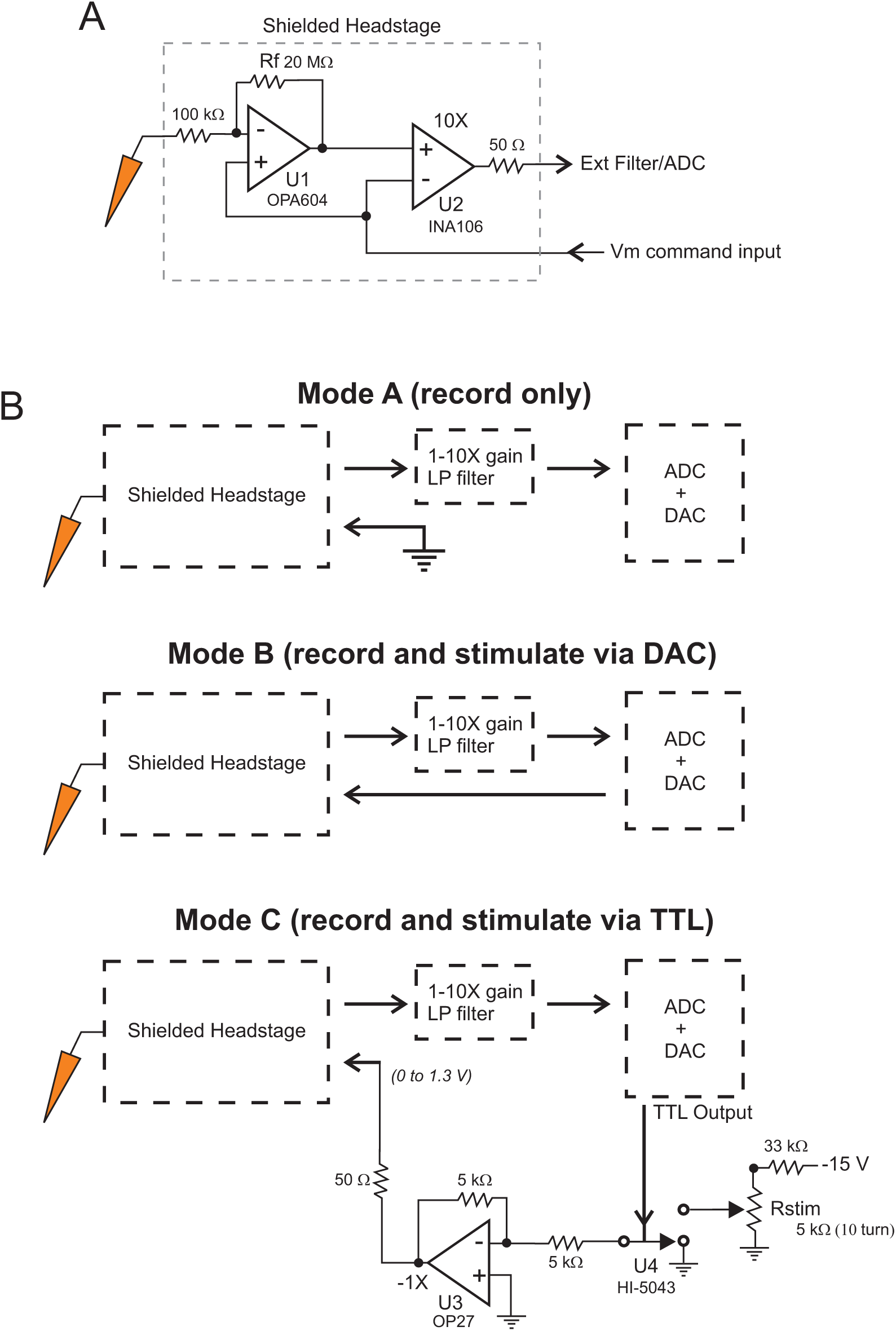
Loose cell-attached amplifier circuit diagrams. A, Diagram of the core headstage amplifier used for LCA recording and stimulation experiments. The input to the amplifier is connected to polycarbonate electrode holder (red). The headstage circuitry is implemented on a custom printed circuit board and mounted inside a small Pomona aluminum box attached to a micromanipulator. The input and output of headstage circuit are connected via thin coaxial cables (RG174) soldered directly to the printed circuit board. All resistors are standard 1% 0.25 watt values. The circuit also included 0.1 *μ*F ceramic bypass capacitors on the *±* 15 V power supply lines to each op amp (not included in the circuit diagram in A). B, Diagrams of different recording modes possible using the headstage circuit shown in A. In *Mode A*, the command input is grounded, allowing a recording-only configuration. The headstage output is typically amplified by 1-10X and low-pass filtered at 10-20 kHz using a Cygnus Technology FLA-01 filter. In the experiments presented here,the ADC+DAC data acquisition system was an Instrutech ITC-18. In *Mode B*, The DAC output of the data acquisition system is used to generate the command signal to the headstage circuit. In *Mode C*, the data acquisition system only provides a timing signal for the stimu-lation pulses. Stimulus intensity was controlled by a 10-turn potentiometer connected to an analog switch (U4) and buffered by a low noise op amp (U3). Buffering the command input using this circuit enables additional command inputs (not shown) to be combined into the voltage signal driving the headstage amplifier.

The patch clamp electrode was connected through a removable polycarbonate holder with a 2 mm pin (A-M Systems #672443). A standard 2 mm female test pin jack (Digikey #J117-ND) was mounted on the Pomona case. The tight fit between the test pin and jack provided sufficient mechanical stability for repeated loose patch recordings provided the electrode holder was seated firmly on the jack. The voltage command input to the head-stage circuit was connected via a RG174 cable. The output of the circuit was connected to another RG174 coaxial cable after passing through a 50 Ω impedance-matching resistor.Both input and output RG174 cables were soldered directly to the headstage printed cir-cuit board with male BNC jacks mounted on the other end of the cables. Design files for the headstage printed circuit board are available at the StrowbridgeLab repository on GitHub (*https://github.com/StrowbridgeLab/LPA*).

While the printed circuit board is small enough to fit inside a miniature Pomona box (Pomona #2400 or #2428), in practice it was more convenient to mount the headstage circuit board inside a longer box (Pomoma #5255) so that the electrode input test pin jack did not have to be placed directly above the board. The longer enclosure also enabled the electrode holders to be located at approximately the same position on the end faces of both the AxoPatch 1D and LCA amplifier headstage boxes, facilitating experiments in which neurons initially recorded using the LCA headstage are subsequently re-patched to establish a whole-cell recording configuration. The headstage box was mounted on a motorized 3-dimensional manipulator (461XYZ, Newport) controlled by custom servo power amplifiers. Custom software enabled a precision computer gaming joystick (Extreme 3D Pro, Logitech) to move the electrode to form the loose patch recording configuration under IR-DIC visu-alization (typically through a Zeiss 63x water objective and imaged with a 0.5 inch CCDcamera onto an analog monochrome monitor). The headstage amplifier was powered by ±15 volt supplies using a regulated linear triple output power supply module (HBAA-40W-A, Power-One). The positive 5 volt output line was not used in the headstage circuit but is required for the accessory analog gating module described below.

### Circuit operation

The headstage circuit can be operated in three different voltage-clamp modes, two of which require only an additional low-pass electronic filter and data acquisition interface to form a complete system. For recording-only applications (*Mode A* in Fig. 1B), the voltage command input to the headstage circuit is grounded and the output signal is digitized after passing through a Bessel low-pass filter (FLA-01, Cygnus Technologies). The circuit ground is connected to the bath ground wire. An example of this recording mode is shown in Fig. 2 from an experiment in which one olfactory bulb mitral cell was simultaneously recorded under the whole-cell (with an AxoPatch 1D amplifier) and loose patch configuration. The neu-ron was stimulated by injecting a depolarizing current step through the whole cell electrode and the resulting train of APs were easily identified in both whole-cell and LCA amplifier outputs. Since the LCA amplifier was not used to stimulate the neuron in this experiment, no artifact subtraction was required in the software routines used to display the LCA output. The LCA recording pipette contained 124 mM NaCl, 3 mM KCl and 10 mM HEPES (pH 7.4).Continuous suction (0.25 - 0.4 PSI; 1700 - 2800 Pa) was applied to the pipette throughout the experiment. Stronger suction was applied when first acquiring the LCA recording configuration. The electrode geometry was similar to those used for conventional whole-cell recording from small interneurons.

**Figure 2:**
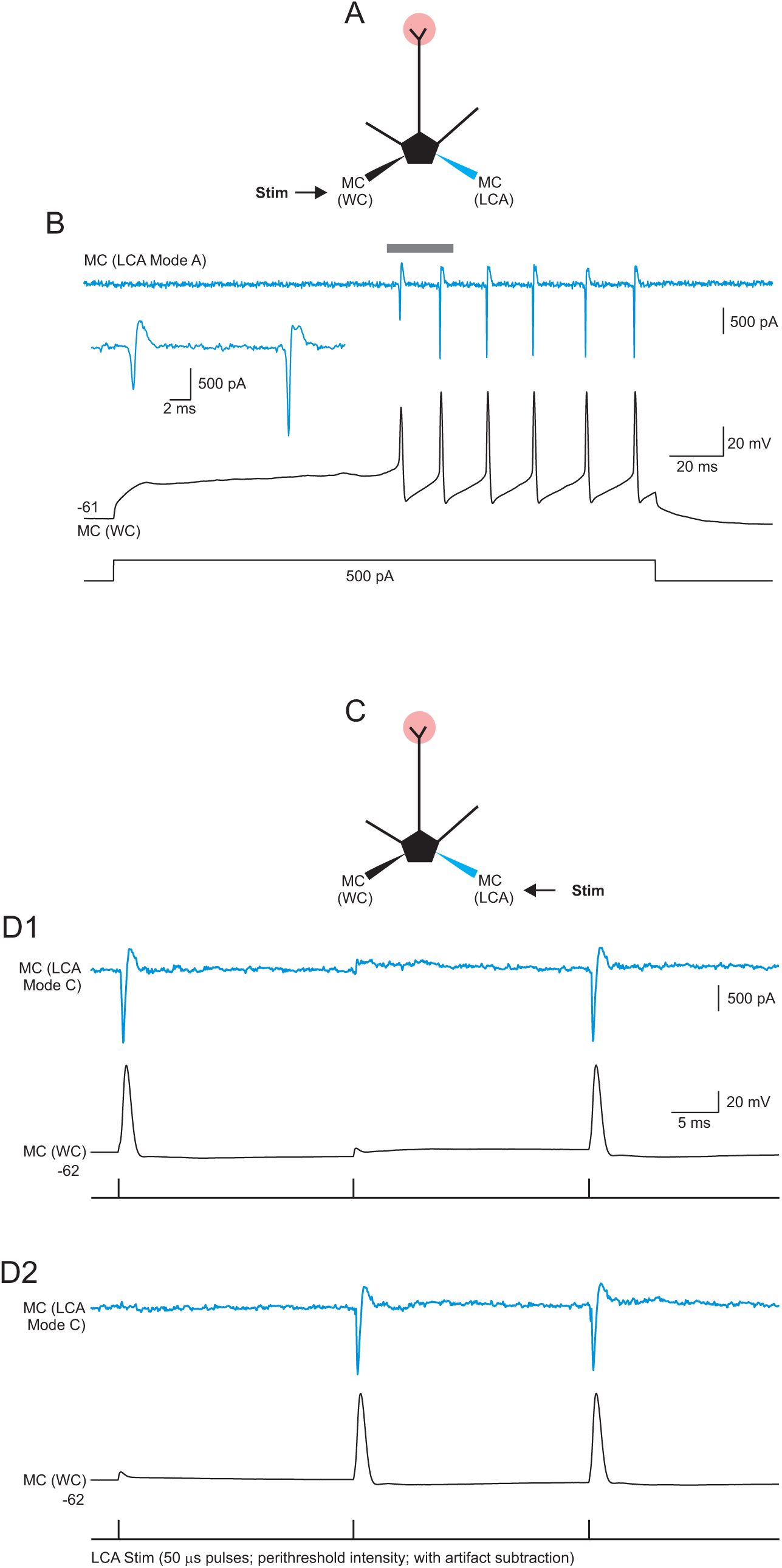
Simultaneous intracellular and loose cell-attached recordings. A, Diagram of the *Mode A* recording configuration. B, Simultaneous whole-cell (black trace) and LCA (blue trace) recordings of the response of a mitral cell (MC) to a depolarizing current step injected through the whole-cell (WC) recording electrode. Inset shows an enlargement of the first two APs recorded in the LCA system. C, Diagram of the *Mode C* recording configuration. D1-D2, Examples of simultaneous intracellular (black traces) and LCA recordings(blue traces) of responses in the mitral cell to brief (50 *μ*s duration) voltage pulses applied through the LCA pipette. The LCA command voltage was set to the perithreshold intensity so that the same stimulus elicited both single APs and failures. (The middle response in D1 and the first response in D2 were failures.) The stimulus artifact (recorded from a single failure trial) were subtracted from each response.

The other two recording modes allow for simultaneous recording and stimulation of the recorded neuron through the LCA. While it is possible to directly control the stimulus voltage applied to the loose patch using the DAC output of the data acquisition interface (*Mode B* in Fig. 1B), this approach requires frequent interaction with a graphical user interface to adjust the stimulus intensity throughout each experiment. In practice, using an analog switch and a 10-turn potentiometer to control the stimulus voltage was preferable (*Mode C* in Fig. 1B). In this mode, a digital output line from the data acquisition interface controlled a single pole, double throw analog switch (HI-5043, Intersil). The output of the analog switch was buffered using a low noise op amp (OP27, Analog Devices) and connected to the headstage input through a coaxial cable. Since the OP27 amplifier is operated in an inverting mode, a negative voltage is applied from the analog switch to generate a positive stimulus at the loose patch. Depending on stimulus voltage required, a fixed resistor can be connected in series with the potentiometer resistor to decrease the sensitivity. Adding a 33 kΩ resistor in series with a 5 kΩ 10-turn potentiometer decreases the sensitivity from 1.5 V/turn to 133 mV/turn with a 15 V supply. This stimulation gain setting for *Mode C* LCA recordings gave satisfactory results with typical experiments requiring 200 - 500 mV pulses (50 *μ*s duration). While shorter stimulus pulses (25 *μ*s duration) provided better separation between the artifact and evoked AP, the stimulus intensities required to reliably trigger APs were more variable than with slightly longer pulses; 50 *μ*s duration pulses provided a good compromise in our experiments. The TTL output lines on the the data acquisition device used in these experiments (ITC-18, Instrutech/Heka/Harvard Bioscience) were optically isolated. If another, non-isolated data acquisition device is employed optical isolation should be added to the analog switch circuit. The stimulus control circuit was assembled on a generic prototyping printed circuit board and powered by the same triple output linear power supply module as the headstage amplifier.

In Figure 2, the same olfactory bulb mitral cell was simultaneously recorded under whole-cell conditions (with an AxoPatch 1D amplifier) and the custom LCA amplifier operating in *Mode A*. A series of APs were evoked by injecting a depolarizing current step through the WC electrode. The resulting spikes are evident in both WC and LCA amplifier outputs. Examples of combined recording and stimulation through the LCA *Mode C* system is shown in Fig. 2C-D and Fig. 3. Without artifact subtraction, the triggered AP was often detectable by a slight prolongation of the response. However, in many experiments, evoked APs were not evident until a failure trial that did not trigger an AP (or slightly subthreshold response) was subtracted from each response. Figure 2D illustrates peristimulus threshold responses (a failure and a success evoked by the same stimulus and acquired in the same episode) with artifact subtraction. Automatic artifact subtraction was implement using custom Python software routines in which a specific stimulus response could be identified using a movable cursor and stored in a memory buffer. Dual recording/stimulation LCA experiments used the same conditions as recording-only experiments (saline-filled pipettes). A potential complication when performing artifact subtraction is saturation of the LCA output signal, resulting in distorted artifact responses. While the FLA-01 used to low-pass filter the head-stage output can provide additional gain, we typically used the filter at either the unity gain setting or minimal additional gain (e.g., 2X) to prevent saturation at ± 10 V full-scale ADC input range using with the ITC-18 acquisition device.

**Figure 3:**
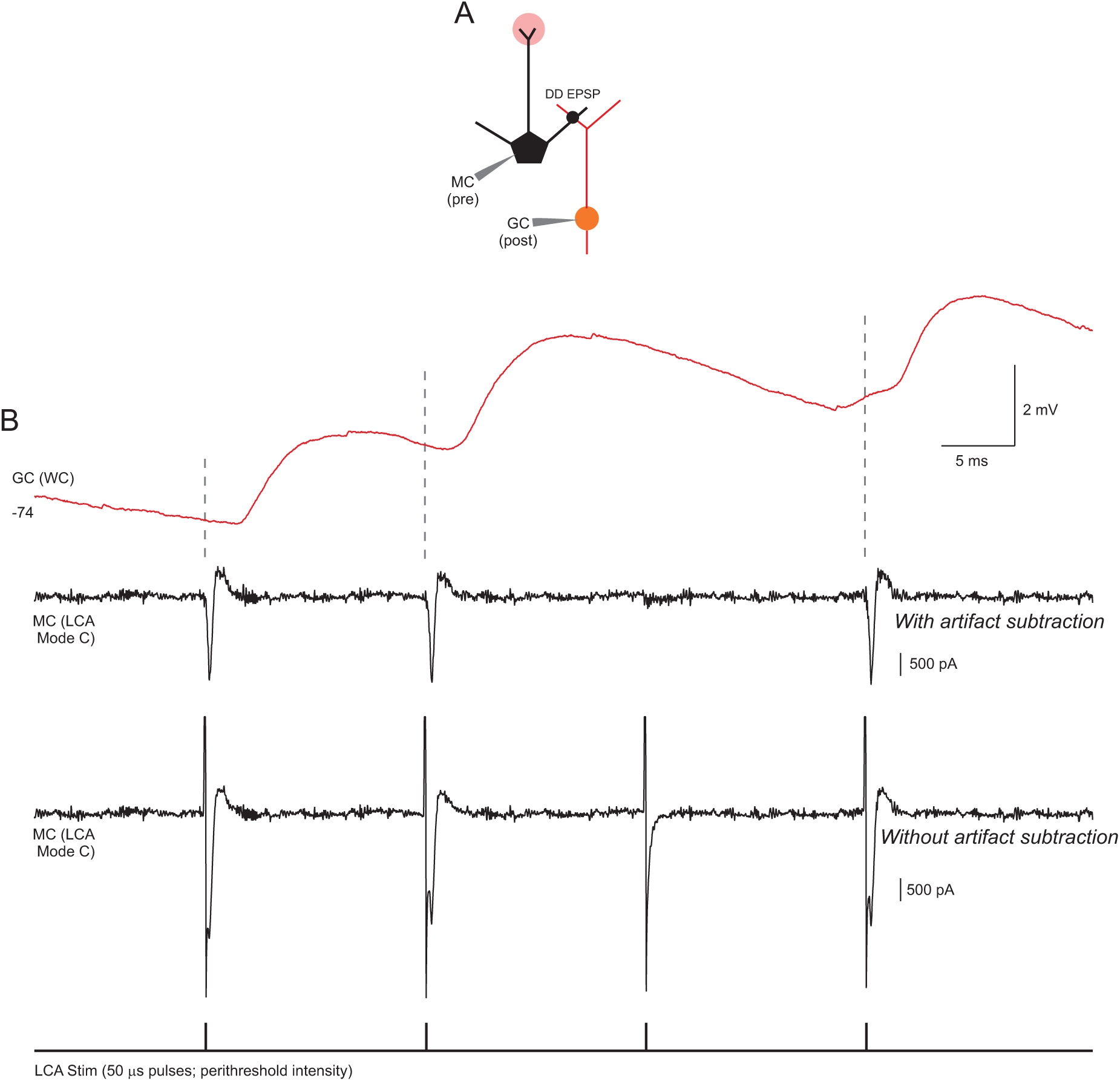
Paired recording between olfactory bulb mitral and granule cells. A, Diagram of recording configuration with a *Mode C* recording from the mitral cell (black traces) and an intracellular (whole-cell current clamp) recording of the granule cell (red trace). Four voltage pulses were applied through the LCA electrode at perithreshold intensity which triggered three APs in the MC (the response to the third pulse was a failure). Each of the MCAPs triggered an EPSP response in the granule at approximately the same latency (1.8 2ms). LCA recording from the MC shown both with artifact subtraction (top black trace) and without artifact subtraction (bottom black trace).

Loose patch clamp recording is ideally suited for testing multiple potential presynaptic neurons that could be synaptically coupled with a postsynaptic target neuron (**5**, **16**). In these experiments, the postsynaptic neuron is typically recorded intracellularly in ei-ther voltageor current-clamp modes to facilitate detection of small postsynaptic responses. An example of a combined LCA/WC paired recording is shown in Fig. 3A-B. In this experiment using olfactory bulb slices, a presynaptic mitral cell is recorded and stimulated using the LCA amplifier in *Mode C* while simultaneously recording from a monosynaptically-coupled granule cell in the current-clamp configuration using an AxoPatch 1D amplifier. At perithreshold LCA stimulus intensities, subthreshold trials that failed to trigger an AP also failed to trigger an excitatory postsynaptic potential (EPSP). Each AP triggered evoked in the mitral cell triggered an EPSP in the granule cell at approximately the same latency (1.8 2 ms) in this example recording.

## Discussion

In this report, we detail the design and operation of a relatively simple amplifier circuit that enables combined recording and stimulation under the loose patch clamp recording configuration. Loose patch stimulation represents a simple method for stimulating one neuron without activating nearby neurons or axons. Loose patch clamp recordings also do not dialyze the intracellular contents of the recorded neuron, providing another advantage over conventional whole-cell recordings. While the relatively large (100s of mV) voltages required to reliably trigger APs is beyond the capacity of most commercial patch clamp amplifiers designed for whole-cell or single-channel recordings, the home-built amplifier we employed uses only two integrated circuits and is inexpensive to construct. The electronic components required for the headstage circuit can be purchased for less than $25 and all the design files required to have printed circuit boards manufactured are available on the GitHub repository. The availability of a simple but workable design for a LCA amplifier that can be used to stimulate APs in individual neurons may be helpful in studies aimed at identifying monosynaptically-coupled neuronal pairs in cell cultures or brain slices. This method could also be applied in vivo to assay the effect circuit and/or behavioral effects of defined manipulations to individual neurons (e.g., **17**).

Our circuit design represents an adaptation of a circuit presented in a previous report (**4**) and is based on classic patch clamp amplifier designs (**10**, **21**). In the previous LCA report (**4**), additional circuitry was employed to transmit both the current signal and command voltage with low noise from a remote electronics box to the headstage circuit. While we did not compare the operation of both circuits, we demonstrate that our simpler design had sufficient signal-to-noise ratio to detect APs in olfactory bulb mitral cells and therefore is likely to be useful in other neurons with large APs such as hippocampal and neocortical pyramidal cells. The present amplifier also did not include circuit elements that were present in the previous Barbour and Isope design that prevented input signal saturation and that isolated the bath ground by passing the reference electrode through a high impedance op amp input. These three non-essential circuit subsystems (paired transmission lines, anti-saturation control and bath ground isolation) accounted for the majority of the components (6/8 op amps) in the previous design. The need for anti-saturation circuitry is reduced in our design because of the lower feedback resistance used in the transimpedance input amplifier (20 MΩinstead of 50 MΩ).

Searching for monosynaptically-coupled pairs of neurons in brain slices is a common application of patch clamp recording (e.g., **11**, **13**, **14**, **16**, **18**, **20**). With whole-cell recordings, glass electrodes typically cannot be re-used after a recording attempt because of the difficulty in achieving a second gigaohm seal once the positive pressure applied to the electrode is released. The inability to re-use patch clamp electrodes represents a significant hurdle during large-scale studies that employ paired recordings since at least one electrode must be removed for every connection tested. The ability to re-use electrodes is one of the primary advantages of LCA recording. With loose patch clamp recording, multiple potential presynaptic neurons can be tested more rapidly while maintaining a conventional whole-cell intracellular recording on the postsynaptic neuron. In our experience using rodent olfactory bulb slices, not having to replace the presynaptic recording electrode after every attempted paired recording attempt increased throughput by approximately 4 times. The primary trade-off with this approach is the inability to visualize the presynaptic neuron without re-patching it to achieve a whole-cell recording configuration. Another trade-off of LCA recordings is the variable threshold voltage required to trigger APs. By periodically monitoring the stimulus voltage required to trigger spikes, we have main-tained monosynaptically-coupled paired recordings for up to 50 minutes (using LCA for the presynaptic neuron and whole-cell recording for the postsynaptic neuron).

A limitation with loose patch stimulation using the present circuit is that for some experiments, a failure trial needs to be acquired and subtracted from each record to reveal stimulus-evoked APs. In many of recordings, differences in the raw current trace are sufficient to enable the operator to determine if an AP was triggered (see Fig. 3B). In some ex-periments, off-line artifact subtraction was required to unambiguously determine when stimuli were effective in triggering APs. In principle, artifact subtraction could be automated by using a second analog switch (and a second TTL output) that reverses the pulse stimulus polarity. In this approach, the acquisition software could apply a stimulus pulse with the same amplitude but opposite polarity at the end of each episode, then automatically display the artifact-subtracted current waveform.

The threshold voltage required to trigger an AP tended to vary throughout most experiments. Since applying large stimulus voltages through the LCA amplifier can degrade the health of the patched neuron, it was helpful to continuously track the threshold stimulus voltage and employ just-suprathreshold intensity stimuli. Even when adjusting the stimulus voltage to be just-suprathreshold, we occasionally triggered spontaneous spiking in the recorded neuron. In most cases, pausing the stimulation for several minutes abolished the spontaneous spiking.

One possible method to reduce the work required to manually track the threshold stimulus is to apply a weak second voltage pulse to monitor seal resistance. This could be implemented using a second analog switch connected to the same virtual ground summing point on the inverting input to U3. By monitoring seal resistance, one could determine if the change in threshold stimulus voltage required to trigger APs could be predicted by changes in seal resistance. If changes in seal resistance explained the variation in the stimulus voltage required to triggered APs, the acquisition software could provide a continuously updated estimate of the threshold stimulus required. Alternatively, the acquisition software could directly control the stimulus intensity using a digital-to-analog converter or a digital potentiometer.

